# Phage interactions may contribute to the population structure and dynamics of hydrothermal vent microbial symbionts

**DOI:** 10.1101/2025.02.21.639545

**Authors:** Michelle A. Hauer, Katherine M. Klier, Marguerite V. Langwig, Karthik Anantharaman, Roxanne A. Beinart

**Author notes:** **Corresponding author**: Roxanne A. Beinart. **Competing Interests:** The authors declare no competing financial interests.

## Abstract

Deep-sea hydrothermal vent ecosystems are sustained by chemoautotrophic bacteria that symbiotically provide organic matter to their animal hosts through the oxidation of chemical reductants in vent fluids. Hydrothermal vents also support unique viral communities that often exhibit high host-specificity and frequently integrate into host genomes as prophages; however, little is known about the role of viruses in influencing the chemosynthetic symbionts of vent foundation fauna. Here, we present a comprehensive examination of contemporary lysogenic and lytic bacteriophage infections, auxiliary metabolic genes, and CRISPR spacers associated with the intracellular bacterial endosymbionts of snails and mussels at hydrothermal vents in the Lau Basin (Tonga). Our investigation of contemporary phage infection among bacterial symbiont species and across distant vent locations indicated that each symbiont species interacts with different phage species across a large geographic range. However, our analysis of historical phage interactions via assessment of CRISPR spacer content suggested that phages may contribute to strain-level variation within a symbiont species. Surprisingly, prophages were absent from almost all symbiont genomes, suggesting that phage interactions with intracellular symbionts may differ from free-living microbes at vents. Altogether, these findings suggest that species-specific phages play a key role in regulating chemosynthetic symbionts via lytic infections, potentially shaping strain-level diversity and altering the composition and dynamics of symbiont populations.

## Introduction

Hydrothermal vents are remarkable habitats found at the ocean floor, where geothermally heated fluids are released through fissures in the Earth’s crust. These ecosystems are sustained by chemosynthetic bacteria and archaea, which convert inorganic molecules expelled from the venting fluid, like hydrogen sulfide and methane, into organic matter. Chemosynthetic microbes at hydrothermal vents exist as both free-living in the water column, rocks, and sediments, as well as symbiotically with foundation species of animals like tubeworms, mussels, and snails, where they provide their hosts with vital nutrients via chemosynthesis (1). Despite the crucial role of symbiotic bacteria in these ecosystems supporting key animal species, phages—viruses that infect bacteria—remain poorly studied among animal-associated populations of chemosynthetic bacteria at hydrothermal vents. Most existing research focuses on phages associated with free-living microbes, leaving a significant gap in our understanding of phage dynamics in symbiotic microbes. Addressing this issue is essential, as phages could play a key role in shaping symbiont populations and in regulating bacterial-animal symbioses.

Viruses are the most abundant biological entities in the oceans, playing a crucial role in bacterial diversity (2–10), biogeochemical cycling of our oceans (11), microbial food web dynamics (12), and in aiding microbial survival (13,14). Much of what is known about the role of viruses in influencing biogeochemistry and structuring bacterial communities comes specifically from lytic (i.e., virion-producing) phages, which infect and lyse their bacterial hosts. Lysis releases dissolved and particulate material, contributing to biogeochemical cycles and controlling bacterial population dynamics and community composition, for example by preventing any single bacterial strain from dominating an ecosystem (15). Further, microevolution driven by interaction with lytic phages also contributes to strain-level bacterial population structure and variation. In the evolutionary arms race between bacteria and phages, bacteria have evolved CRISPR (<u>c</u>lustered regularly interspaced <u>s</u>hort <u>p</u>alindromic repeats) spacers, viral DNA fragments incorporated into bacterial genomes that serve as an immune memory and enable bacteria to recognize and prevent future invaders (16). These spacers, therefore, represent a historic catalog of some of the past attempted infections of bacterial hosts. Studies have found that phages may play a role in bacterial population structure (17–20) including at the strain-level (21,22), and there is ongoing research in the use of CRISPR as a hypervariable region for micro-evolution (23). In contrast, lysogenic phages directly alter their host bacterium’s genome by inserting their own genetic material into the bacterium’s chromosome, becoming a “prophage” that is replicated through binary fission. Since a prophage requires use of its host bacterium’s machinery for its own replication, it can be advantageous for prophages to enhance their host fitness via auxiliary metabolic genes (AMGs) (13,14) — viral genes originally derived from bacteria that that can be used to modulate the host bacterium’s metabolic activities and contribute to biogeochemical processes, by enhancing a bacterium’s nutritional absorption, survival in unfavorable conditions, and pathogenicity (12,24–31). Altogether, through both ecological and evolutionary processes, interaction with lytic and lysogenic phages can contribute to genomic and phenotypic variation within and among bacterial species.

Temperate phages, which can integrate their genomes into bacterial genomes as prophages and switch between lytic and lysogenic cycles, are notably abundant at hydrothermal vents relative to other marine habitats (32,33), often harboring AMGs that modulate their host bacterium’s metabolism (e.g., nitrogen and sulfur metabolisms) (14,34–36). The high prevalence of prophages at vents are attributed to the geochemical dynamism of these habitats, which facilitates horizontal gene transfer (HGT), resulting in a broader functional gene set that may increase microbial adaptability to the extreme environmental fluctuations (37,38). Although prophages are abundant in free-living microbes at vents (35), not much is known about the abundance, diversity, or role of phages in influencing symbiotic microbes. Nonetheless, interest in the topic has grown rapidly, with recent studies from deep-sea hydrothermal vent symbioses harboring mixed results regarding the presence of prophages and the possibility of an AMG-mediated phage-microbe-animal tripartite symbiosis (37,39–42). Furthermore, recent population genomic studies suggest bacteriophages may also play a role in strain-level population structuring of microbial symbionts from deep-sea hydrothermal vents (21,22), though there is currently a dearth of knowledge regarding the role of prophages or CRISPR spacers in symbiont population structuring (48).

This study investigates phages associated with the chemosynthetic bacterial symbionts of foundation mollusk species at six deep-sea hydrothermal vent fields separated by 10s to 100s of kilometers in the Lau Basin (Tonga) (Figure 1). Specifically, we investigated evidence for interaction with phages in the chemosynthetic symbionts of IUCN Red-List Endangered and Vulnerable (https://www.iucnredlist.org) sympatric snails *Alviniconcha boucheti* (43), *Alviniconcha kojimai* (44), *Alviniconcha strummeri* (45), and *Ifremeria nautilei* (46), as well as the mussel *Bathymodiolus septemdierum* (47). Each host species harbors 1-2 specific intracellular endosymbionts in their gill tissue, with gammaproteobacterial symbionts *Candidatus* Thiodubiliella endoseptemdiera (Family Thioglobaceae) in *B. septemdierum,* gamma1 (Family Thiomicrospiraceae) in *A. kojimai* and *A. strummeri*, GammaLau (Family Sedimenticolaceae) in *A. strummeri*, thiotrophic SOX (Family Arenicellaceae) and methanotrophic MOX (Family Methylomonadaceae) in *I. nautilei*, and campylobacterial symbionts *Sulfurimonas Epsilon* and *Sulfurimonas sp.* in *A. boucheti* (Family Sulfurimonadaceae) (50). Previous research on these and other chemosynthetic symbionts have demonstrated that they commonly exhibit strain-level differentiation by biogeography, with evidence of local adaptation potentially influenced by viral assemblages (21,22) suggesting interaction with phages may be important to their ecology and evolution. Given this, we examined the lytic phage communities present in symbiont-containing tissues and the prophage and CRISPR spacer content in symbiont genomes to assess the potential variation in symbiont-phage interactions by host species and geography.

**Figure 1.**
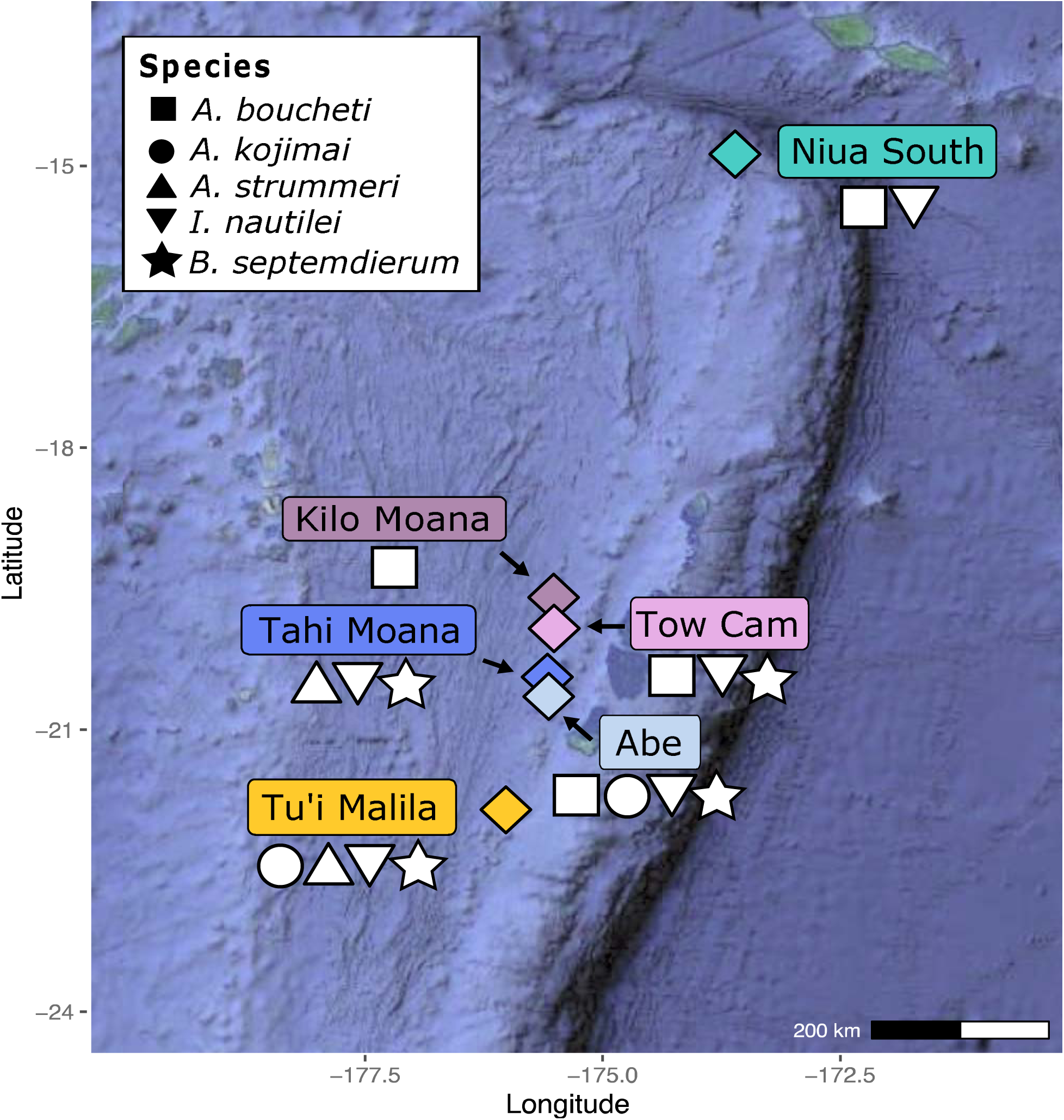
Map of the Lau Basin hydrothermal vent fields from which animal specimens were collected for this study. Vent field color scheme used in this figure is consistent through all downstream figures. Figure adapted from Breusing et al. 2022 (21).

## Methods

### Phage detection and host prediction in symbiont MAGs and animal metagenomes

All 219 metagenomes (Table 1) and 200 symbiont MAGs (Table 2) used in this study were previously sequenced and assembled in Breusing et al., 2022 and 2023 (21,48) Prophage content in symbiont MAGs as well as both lytic and lysogenic phages in metagenomes were identified using VIBRANT v1.2.1 (49). Since non-symbiotic bacterial taxa and their phages could potentially be physically associated with the gill tissues (e.g., on gill surfaces) and, therefore, present in our metagenomes, we predicted whether the lytic phages identified in the animal metagenomes target the expected symbionts using iPHoP v1.3.3 (50) with symbiont MAGs added to the iPHoP database and default parameters. Lytic phages identified as targeting the expected symbionts with ≥90% confidence in the iPHoP output were considered symbiont-targeting phage. If at least one individual phage within a cluster was identified as symbiont-targeting, the entire species cluster was designated as such.

**Table 1.** Total number of metagenomes analyzed, and the average number of symbiont-targeting phage with ≥10kbp and ≥4 ORFs per Mbp of metagenome.

| Host Animal Metagenome | No. Metagenomes | Average no. symbiont-targeting phage per Mbp | No. Species Clusters | No. Symbiont-targeting Species Clusters |
| --- | --- | --- | --- | --- |
| *I. nautili* | 59 | 0.00307 | 19 | 15 |
| *A. boucheti* | 23 | 0.00023 | 3 | 2 |
| *A. kojimai* | 17 | 0.01626 | 8 | 5 |
| *A. strummeri* | 22 | 0.00632 | 9 | 4 |
| *B. septemdierum* | 98 | 0.00475 | 70 | 59 |
| Total | 219 |  |  |  |

**Table 2.** Total number of MAGs mined; total number, percent, and average number per Mbp of lysogenic phage (≥10kbp, ≥4 ORFs) found in each MAG as identified by VIBRANT; and total and average number per Mbp of CRISPR spacers found per symbiont type.

| Symbiont MAG | No. MAGs | No. prophage | % MAGs with lysogenic phage | Average no. prophage/Mbp | Total no. spacers | Average no. Spacers/Mbp |
| --- | --- | --- | --- | --- | --- | --- |
| SOX - *I. nautiliei* | 58 | 0 | 0 | 0 | 1157 | 4.5653 |
| MOX - *I. nautiliei* | 3 | 0 | 0 | 0 | 0 | 0.0000 |
| Epsilon - *A. boucheti* | 8 | 0 | 0 | 0 | 32 | 2.3158 |
| *Sulfurimonas* - *A. boucheti* | 6 | 0 | 0 | 0 | 47 | 1.5991 |
| Gamma 1 - *A. strummeri* | 19 | 4 | 21% | 0.036 | 21 | 0.2957 |
| Gamma1 - *A. kojimai* | 16 | 14 | 88% | 0.203 | 215 | 3.4597 |
| *Ca. T endoseptemdiera* - *B. septemdierum* | 92 | 0 | 0 | 0 | 1920 | 7.6743 |

### Dereplication and taxonomic classification of bacteriophages

To create species-level phage clusters to assess phage infection patterns, phage nucleotide sequences produced by VIBRANT were dereplicated with dRep v3.4.5 (51) using 95% Average Nucleotide Identity (ANI) and 50% minimum alignment for both lytic and lysogenic phages from the metagenomes and prophages from the MAGs. Phage sequences ≥10kbp and ≥4 open reading frames (ORFs), as filtered by dRep, were retained for downstream analyses. To visualize the phage clusters, we used the cluster dendrogram generated with dRep. We circularized the dendrogram by modifying the radialtree code and integrated it into the dRep code (Original radialtree: https://github.com/koonimaru/radialtree/blob/main/radialtree.py, Original dRep: https://github.com/MrOlm/drep/blob/master/drep/d_analyze.py, forked files in https://github.com/michellehauer/Lau_Basin_09-16_Phage_Analysis/blob/main/drep_fork/) using Python v3.8.9. (52). The relative frequency of each symbiont-targeting phage species cluster was calculated as the proportion of phages in a given cluster found at a given location relative to the total number of phages in that cluster and was represented in the bubble heatmap figure using ggplot2 (52) in RStudio v2023.12.1+402. Phage taxonomic classification was performed using geNomad v1.6.1 (53) using default parameters, and only classifications with ≥5 gene alignment and an agreement of ≥80% were considered valid.

### Investigation of genetic elements contributing to symbiotic functionality in symbiont MAGs

We screened prophages for AMGs using VIBRANT (49) and only retained AMGs flanked by viral genes, viral-like genes, or hypothetical genes of likely viral origin (based on V-scores). For AMGs detected in an individual prophage that belonged to a dRep species cluster, a blastp v2.101 analysis was performed to query the AMG protein sequence against a database of all prophage protein sequences from that prophage species cluster, followed by a mafft v7.505 alignment.

### CRISPR Spacers in Symbiont MAGs

CRISPR spacer abundance and diversity were determined by MinCED v0.4.2 using default parameters (https://github.com/ctSkennerton/minced). Spacer taxonomic classifications were determined by a blastn analysis with CRISPR spacer sequences against the nt virus database using blast-plus v2.13.0 and Python 3.11.6 (54). We defined a CRISPR spacer “match” as having 100% identity across ≥20bp, parameters shown to be suitable for determining the species or genus from which the spacer derived (16). Plasmid sequences were removed by screening spacers against the PLSDB (Plasmid Database, v2023_11_03_v2) (55) using MASH Screen and default parameters of max. p-value 0.1 and percent identity 99% on the web user interface. To determine which form a phage species cluster, Cd-Hit v4.8.1 (56) was used with ≥85% sequence similarity. For each CRISPR spacer cluster, the distribution across locations was determined by calculating the proportion of spacers found at each location relative to the total number of spacers in that cluster. This proportion was then converted to a percentage to illustrate the relative presence of each CRISPR spacer cluster at different locations using a heatmap created with ggplot2 (52) and pheatmap (57) in RStudio v2023.12.1+402. To determine whether symbionts harbor spacers that match phages found in metagenomes, a reciprocal blast analysis with parameters-word_size 20 and -max_target_seqs 1 (54) was performed using the “best hit” phage per species cluster as determined by dRep. Results with ≥97% percent identity and ≤1 mismatch were considered a match.

## Results

### Symbiont-containing animal gill tissues harbor distinct bacteriophage viromes that do not exhibit strict geographic endemism

A total of 225 individual phages, averaging 0.0059 phage per megabase pair (Mbp), were identified across the 219 metagenomic samples (Figure 2, Supplementary Table 1-2), 81.53% of which were predicted to target the symbionts (Figure 2, Supplementary Table 3-5). A total of 105 species-level phage clusters were formed across all phages in the metagenomes, though 84 (80%) comprised of a single, unique phage sequence detected only in one snail or mussel’s metagenome and did not form a species-level cluster with any other sequence (Figure 2, Supplementary Table 3-4). Of these 105 phage species clusters, 91 were comprised of symbiont-targeting phage species (Figure 2, Supplementary Table 5).

**Figure 2.**
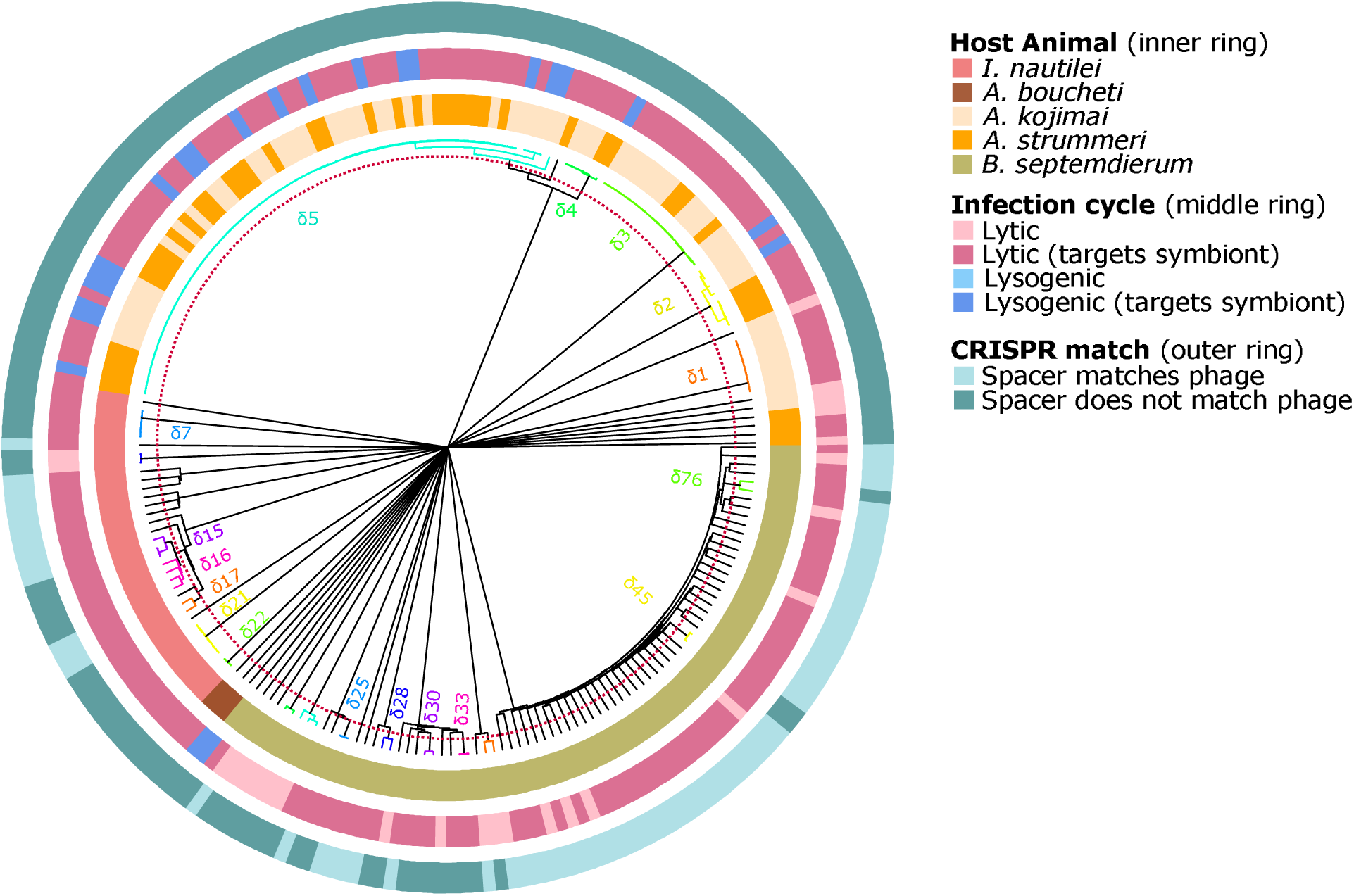
Circle dendrogram representing all lytic and lysogenic phage collected across all metagenomes. Each branch corresponds to a phage, with species clusters defined by 95% ANI, indicated by the red dotted line. These species clusters are color-coded for enhanced clarity, and black branches denote singletons. dRep does not calculate or visually represent branch relationships for sequences with less than 88% ANI. The innermost ring illustrates the source animal metagenome from which the phage was derived. The middle ring represents the phage’s cycle and indicates whether it was predicted to target the symbiont; if at least one individual within a cluster was identified as symbiont-targeting, the entire cluster was designated as such. The outermost ring is a binary ring that signifies that a CRISPR spacer from one of the symbiont MAGs matched to that phage sequence.

Except for four phage species clusters present in both the *A. kojimai* and *A. strummeri* metagenomes, most phage species clusters were uniquely observed in the metagenome of only one host animal species, suggesting distinct bacteriophage viromes infect their respective microbiomes (Figure 2, Supplementary Table 3). Four phage species clusters (φ2-φ5) were shared between *A. kojimai* and *A. strummeri*, though only φ3, φ4 and φ5 targeted the gamma1 symbiont (Supplementary Table 3, 5). Phage species clusters φ4 and φ5 were both predicted to contain a mix of lysogenic and lytic forms (Figure 2, Supplementary Table 1), whereas φ3 was exclusively lytic.

Across all host animal metagenomes, phage species did not consistently exhibit strict geographic endemism at vents within the Lau Basin (Figure 3, Supplementary Table 3). Of the 18 total symbiont-targeting phage species clusters that were identified in more than one individual metagenome, 10 were restricted to a single vent field (e.g., φ1 in *A. kojimai,* φ4 in *A. kojimai* and *A. strummeri* metagenomes, φ15 and φ22in *I. nautilei*, and φ25, φ30, φ33, φ37, φ45, φ76 in *B. septemdierum*). The other 8 were associated with the same host species across two or more Lau Basin vent fields.

**Figure 3.**
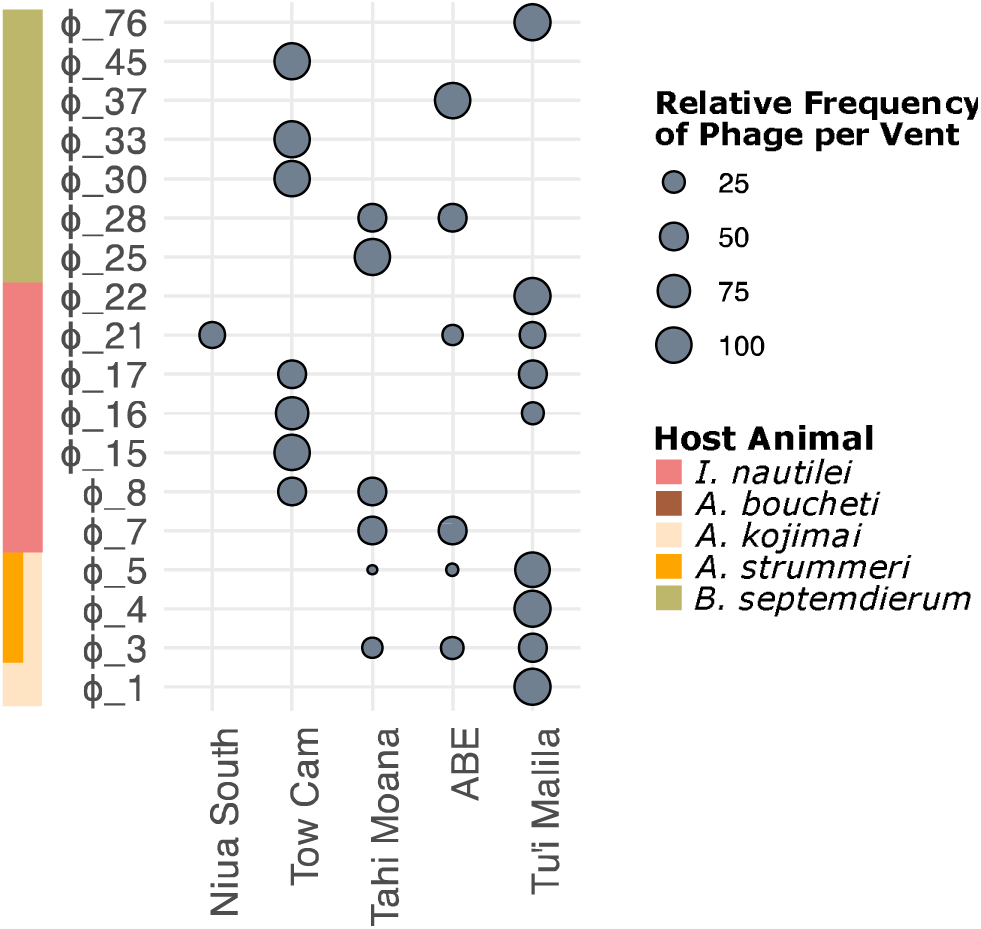
Bubble heatmap representing the proportional distribution of the occurrence of each phage species cluster in the metagenomes by hydrothermal vent field (ordered north to south). Only the phage species that were detected in more than one individual metagenome were included (i.e., excludes singletons).

### Prophages found exclusively in gamma1 symbionts

Prophages were only detected in the gamma1 symbiont MAGs of *A. kojimai* and *A. strummeri* (Figure 2, Supplementary Figure 2, Supplementary Table 6-8); no other symbiont species harbored prophages. Among the gamma1 symbionts from *A. kojimai* and *A. strummeri,* 12/16 (75%) and 4/19 (21%) harbored one or more prophage, representing an average of 0.19 and 0.049 prophage/Mbp, respectively (Supplementary Table 6).

Of the gamma1-*A. kojimai* prophages, 10 were found in individuals from Tu’i Malila and 2 at ABE; all 4 gamma1-*A. strummeri* MAGs were found at Tu’i Malila (Figure 3, Supplementary Figure 1b-c). These prophages formed two distinct species clusters (95% ANI)— φ4 and φ5—but three distinct strains (99% ANI) (Supplementary Figure 2, Supplementary Table 8-9). Species cluster φ4 was found in a single *A. strummeri*–gamma1 MAG from Tu’i Malila, and the remaining 15 prophages belong to species cluster φ5. Within φ5, gamma1-*A. kojimai* gamma1 symbionts from ABE formed a distinct strain-level group, while both *A. strummeri* and *A. kojimai* gamma1 prophages from Tu’i Malila formed another distinct strain. These results may indicate that prophage species, like the phages in the metagenomes, do not exhibit strict species-level geographic endemism; however, they may segregate at the strain-level by vent field.

### Phages predominantly belong to the class *Caudoviricetes*

Of the taxonomically classifiable phages and CRISPR spacers, most species were from the dsDNA class *Caudoviricetes* (Supplementary Table 3,7,10), a common and highly diverse marine phage family previously reported to be abundant at hydrothermal vents (36). Three lytic *Caudoviricetes* phage clusters from Tow Cam—φ30, φ32, and φ34—were classified within the order *Schitoviridae*. Only one individual lytic phage—φ104 at Tu’i Malila—was of the class *Malgrandaviricetes* in the order *Microviridae*, a ssDNA bacteriophage also commonly found in hydrothermal vent ecosystems including the Lau Basin (34,36).

### Limited AMG Occurrence in Symbiont-Infecting Prophages

One gamma1 prophage (species φ5) from *A. kojimai* (K09_101 at ABE) was identified to harbor an AMG. This AMG was annotated as glutamine---fructose-6-phosphate transaminase (*glmS*) (Supplementary Table 11), a gene that is involved in UDP-GlcNAc synthesis, a precursor of peptidoglycan for bacterial cell wall formation, which is not clearly auxiliary. Furthermore, the BLASTp analysis of the *glmS* gene from K09_101 did not have any reliable matches or alignments to the other protein sequences from individual phages in species cluster φ5, providing no substantial evidence for widespread occurrence of this *glmS* gene among the chemosynthetic symbionts examined here.

### CRISPR spacer clusters exhibit high host-specificity

Among all MAGs, 58% contained at least one CRISPR spacer, with counts ranging from 1 to 230 per MAG (Supplementary Table 12), and no spacers were identified as plasmid derived. No spacers were identified in *I. nautilei* MOX symbionts, whereas *Ca.* T. endoseptemdiera (*B. septemdierum*) symbionts harbored the greatest average number of CRISPR spacers per Mbp and gamma1 symbionts had the lowest average number of spacers per Mbp (Table 2, Supplementary Table 12).

CRISPR spacers found in the symbiont MAGs formed 2,679 clusters, 544 (20%) of which harbored more than one spacer (i.e., were not singletons) (Supplementary Table 13). The remaining 2,135 (80%) CRISPR spacers were singletons (only occurred in one MAG), with the exception of three spacer clusters in *A. strummeri* and *A. kojimai* gamma1 MAGs: 128, 154, and 179 (Figure 4, Supplementary Table 12). Thus, CRISPR spacers were largely segregated by symbiont species and, unlike prophage infections, the majority of CRISPR spacers in gamma1 symbionts exhibited host-specificity at the strain level (Figure 4, Supplementary Table 13). Like the phage species clusters from the metagenomes, some of the CRISPR spacers did not exhibit strict geographic endemism (Figure 4), though many spacers were indeed unique to a given vent field, with endemism particularly more common at Tow Cam.

**Figure 4.**
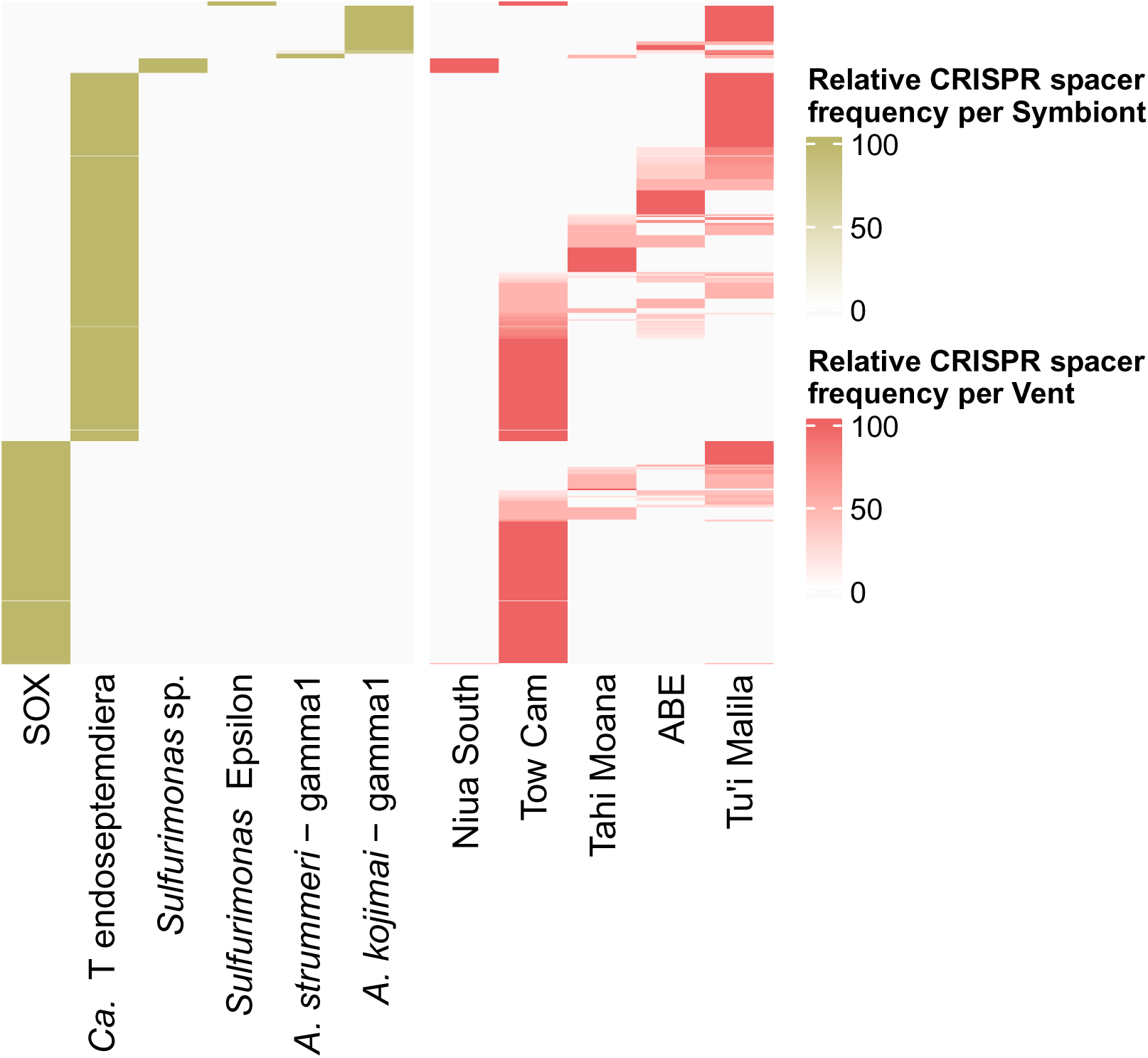
Heatmap of relative abundance of the occurrence of CRISPR spacers clusters in the symbiont MAGs by phylotype and vent field. Y-axis represents the spacer clusters. Only the CRISPR spacers that were detected in more than one MAG were included. Vent fields are ordered from north to south. (53)(57)

### CRISPR spacers indicate active immune system mechanisms in SOX and *Ca.* T. endoseptemdiera

Some CRISPR spacers found in the SOX symbionts of *I. nautilei* and the *Ca.* T. endoseptemdiera symbionts of *B. septemdierum* matched symbiont-targeting lytic phage sequences that were present in the metagenome, therefore, possibly representing active immune systems (Figure 2, Supplementary Table 14). Specifically, 66 lytic phage-spacer matches were found in SOX and 280 lytic phage-spacer matches were found in the *Ca.* T. endoseptemdiera symbiont MAGs. In some cases, many CRISPR spacers (up to 27) matched to a single phage sequence.

## Discussion

Strain-level variation among microbial symbionts can significantly influence the ecology and evolution of their host animals (58), yet our understanding of what drives the strain-level variation remains limited, especially as it pertains to the role of bacteriophage. Here, we investigated the communities of lytic phage populations associated with symbiont-containing gill tissues of five co-occurring endangered and vulnerable deep-sea hydrothermal snail or mussel species from hydrothermal vents in the Lau Basin, as well as the integrated prophages and CRISPR spacers in their symbiont MAGs. In summary, our results indicated that lytic phages present in the gill tissues primarily targeted the dominant bacterial symbionts in that tissue, and that prophages and associated AMGs were not detected in most symbiont species. Overall, symbiont-targeting lytic and lysogenic phages exhibited host-specificity at the species (but not strain) level, and did not exhibit strict geographic endemism among hydrothermal vent fields in the Lau Basin. In contrast, CRISPR spacers in symbionts largely demonstrated host-specificity, including at the strain-level. Although many spacers lacked endemism, a considerable number were nonetheless unique to specific vent fields, highlighting both species-specific and biogeographic influences on spacer distribution and their possible role in shaping symbiont population structure.

Whereas some phages are generalists and cosmopolitan, most have high host-specificity at the species-level (59) or even at the strain-level (60–63), and vent-dwelling phages have typically been found to be endemic (34,64). The chemosynthetic symbionts of mollusks from the Lau Basin exhibit vent-based geographic population structure that appears to be driven in part by phage interactions (21,22); we, therefore, expected to observe unique vent-endemic phage infections and CRISPR spacers among our symbiont strains based on geography. However, neither the phages nor CRISPR spacers in this study exhibited consistent, strict geographic endemism. It is generally thought that patterns of biogeographic structuring in phages are due to environmental selection and host availability (65), as opposed to physical barriers limiting phage dispersal (65). Given the wide distribution of host availability among vent locales and the limited physical barriers to dispersal in the Lau Basin (21), the lack of phage biogeographic structuring or strict endemism agrees with many other models of phage distribution. However, whereas some of the phage species clusters were not geographically endemic, strain-level biogeographic structuring was observed in phage species cluster φ5 within gamma1 symbiont MAGs, suggesting localized adaptation. Furthermore, the absence of biogeographic structuring in some phage clusters might be attributable to the proximity of the ABE and Tahi Moana vent fields, which are only 8 km apart. Previous work demonstrated that the symbionts show strain-level variation by biogeography *except* for at ABE and Tahi Moana (21), suggesting that these vent fields could perhaps be treated as one geographic locale. Nonetheless, many phage species are shared between the two southernmost vent fields, ABE and Tu’i Malila, which are ∼141km apart. In some cases, phages span even greater distances, such as cluster φ21, where viruses of *I. nautilei* SOX were found at the northernmost vent field Niua South as well as the southernmost two vent fields, ABE and Tu’i Malila, which are ∼821km apart. Although phage infections of SOX and gamma1 were predominantly non-endemic, five of the six phage species clusters infecting *Ca.* T endoseptemdierum symbionts of *B. septemdierum* (φ25, φ30, φ33, φ45, φ76), did indeed appear to be endemic. One exception, φ28, was found at both Tahi Moana and ABE. These results suggest that contemporary phage infections may play a stronger role in structuring the biogeographic strain-level variation among *Ca*. T endoseptemdierum symbionts relative to the other chemosynthetic symbionts within the Lau Basin.

Given that these symbiotic mollusks typically harbor one or two dominant bacterial symbiont species at high cell densities in their gill tissues, comprising the majority of their gill microbiome, it is unsurprising that most of the lytic phages in the tissue samples were predicted to infect the symbionts. These symbionts predominate the gill tissue microbiomes, and although their abundance and density has not been directly assessed, a similar vent mussel species, *Bathymodiolus puteoserpentis,* has been estimated to harbor a mean abundance of 2.5×10^12^ chemosynthetic bacterial symbionts per specimen (66). Whereas most phage species were found exclusively in one host animal metagenome, *A. strummeri* and *A. kojimai* had more similar bacteriophages, and these were largely predicted to infect gamma1 symbionts, the bacterial symbiont that these snail species share (67). This finding suggests that animals typically harbor distinct phage viromes, underscoring the specificity of these phages (59–63).

The prophages and CRISPR spacers of bacterial symbiont MAGs exhibited species-level patterns, suggesting that each symbiont species harbors unique viral defenses and prophage associations. Since previous studies suggest that phage interactions contribute to the strain-level variation of bacterial symbionts at the Lau Basin (21,22), and some global viruses are reported to harbor high strain-level host-specificity (60–63), we hypothesized that the CRISPR spacer and prophage infections in the gamma1 symbionts might be strain-specific, having reflected divergent histories of interaction with phage. The vast majority of CRISPR spacers in the gamma1 symbionts were indeed strain-specific based on the host animal from which they derived, whereas the prophages in gamma1 symbionts of *A. kojimai* and *A. strummeri* and appeared to be strain-agnostic. The CRISPR spacer results suggest that historical phage infections had strain-level host-specificity and/or that host-animal specific gamma1 strains maintain distinct CRISPR spacers due to differential fitness consequences from maintaining spacers (67–69). In either case, the results suggest that CRISPR spacers may directly or indirectly contribute to strain-level variation between *A. kojimai* and *A. strummeri*. Although strict geographic endemism was not observed among the spacers, many spacers were indeed unique to a given vent field—in particular at Tow Cam—suggesting possible contribution to biogeographic strain-level variation among the symbionts as well.

Furthermore, many of the CRISPR spacers matched lytic phage sequences in *I. nautilei* and *B. septemdierum* metagenomes, including those that were predicted to infect their respective symbionts, suggesting that not all the CRISPR spacers are necessarily representative of historic infections but may also reflect contemporary interactions. In contrast, the gamma1 spacers did not match to any active infections, and, therefore, gamma1 spacers likely only represent historic infections. Additionally, the relatively low number of CRISPR spacers found in gamma1 compared to the other symbiont species may suggest that their CRISPR-cas immune system is less active. Indeed, the gamma1 symbionts were the only species in this study to harbor detectable prophages, but also had the lowest average number of CRISPR spacers. Whether this lack of spacer protection is an evolutionary advantage, since it may more readily allow for the integration of beneficial prophages, is unknown. If a prophage infection is indeed advantageous for gamma1 symbionts, whether through AMGs, horizontal gene transfer (HGT) (70,71), or super immunity (72,73), an active CRISPR immune system may present a fitness cost. Instead, gamma1 symbionts may prefer to employ other mechanisms to prevent infection from unwanted invaders, e.g., via restriction modification genes or directly through the super immunity potentially provided by prophage integration. This may also provide insight into the growth rates of the bacterial symbionts, in that faster growing symbionts have been observed to harbor diminished CRISPR-Cas immunity and greater number of prophage relative to slower growing bacteria (74).

The lack of prophage detection in all other symbionts was unexpected, considering their prevalence in vent bacteria (32,33), and previous findings of prophages in *B. septemdierum* symbionts (42). Our more conservative parameters (requiring phage sequences to be ≥10kbp) likely contributed, as our raw data did indicate a small number of prophages in *Ca.* T. endoseptemdiera at lengths below our criteria of 10kbp. Although this could be an artifact of methodology, other studies of chemosynthetic symbionts also found limited evidence for prophages (40).

The lack of detected prophages in the symbionts might be explained by the fact that intracellular symbionts are sheltered from the extreme chemical and temperature dynamism of hydrothermal vents, whereas free-living environmental microbes may confer beneficial fitness consequences from prophages, as the resulting genetic augmentation could confer adaptive advantages critical for survival in such fluctuating environments (12,24–31). From a game theoretical perspective, the strategy of free-living vent microbes accepting prophages can be seen as a form of genetic hedging (75–77), wherein the potential long-term benefits of acquiring new genes outweigh the costs (e.g., lysogenic induction into the lytic cycle). Conversely, microbes living within host cells face different strategic considerations wherein the relative environmental stability reduces the benefit of such genetic diversity, and the sudden activation of a lytic cycle could lead to rapid and unchecked microbial cell death, disrupting the symbiotic relationship.

Our data provided no clear evidence of prophages being fundamentally involved in the bacterial-animal symbioses at deep-sea hydrothermal vents of the Lau Basin via AMGs. This may be an artifact of the methodology, for example detection limits, genome fragmentation, or the absence of intact lysogenic prophages in symbionts, although prophages with AMGs are more common in habitats where metabolic modification is critical for survival, and, therefore, habitats that are nutrient enriched typically do not harbor prophage with AMGs (72). The overall stability of the host-associated conditions, coupled with likely high nutritional availability, may explain the limited detection of AMGs. Furthermore, not all beneficial prophages are involved in modifying their host’s metabolic pathways; some may primarily provide benefits like increased resistance to superinfection, for example through lysogenic allelopathy (72,73), or horizontal gene transfer (70,71). The temperate phage species clusters in gamma1 symbionts could also act as population control, protecting the host animal from the destabilizing effects of symbiont over-colonization. (79)(79)

## Conclusions

In this study, we investigated the phage content in metagenomes from the symbiont-containing gill tissues of four hydrothermal snail and one mussel species, as well as the prophage and CRISPR spacer content in their bacterial symbiont MAGs, across a ∼821 km range in the Lau Basin. The complex interplay between phages and chemosynthetic symbionts in deep-sea hydrothermal vent ecosystems underscores a multifaceted ecological dynamic that may significantly impact microbial population dynamics. Our study revealed that contemporary interaction with lytic and lysogenic phages do not show a geographic pattern, though we found some evidence for historical infections by strain-specific phage that vary by location. To better understand whether genetic contributions from phages through HGT influence symbiont adaptability and strain-level variation, further research examining symbiont genomes for genes of phage origin (e.g., virulence genes) and expression is warranted. Furthermore, temporal analyses may elucidate the stability of these phage infections and dynamics of symbiont-phage coevolution, providing more clues as to whether some of these phage infections confer fitness consequences. Microscopy of host animal tissue may also provide clarity regarding whether the observed lytic phages in this study are found in the intracellular symbiont populations within the gill tissue or outside on the surface of the gills, which may further influence the interpretation of these interactions.

Given that all five animals in this study are currently classified as “Endangered” or “Vulnerable” on the IUCN Red List (https://www.iucnredlist.org) and play a vital role in the ecology of hydrothermal vents, understanding the factors influencing the success and connectivity of their symbionts is crucial for future conservation and management efforts. Our results indicate that phages may play a multifaceted role in controlling symbiont populations and bacterial-animal symbiotic dynamics, including host death through lytic infection and/or the modulation of strain-specific interactions through CRISPR spacers or prophage integration. These interactions suggest that distinct symbiont strains engage with different phages, contributing to strain-level genomic variation and ecological dynamics. These interactions may thereby impact the ecological stability of these hydrothermal vent communities by preventing overdominance of particular symbiont strains or modulating the fitness of the locally available symbiont strains. Elucidating the role of phages in host-symbiont dynamics may, therefore, be an integral component of informing effective future management strategies.

## Declarations

### Data Availability

The previously published raw metagenomic reads as well as the corresponding MAG assemblies are available at the National Center for Biotechnology Information under BioProject <u>PRJNA855930</u> and <u>PRJNA523619</u>. All bioinformatic scripts are published to https://github.com/michellehauer/Lau_Basin_09-16_Phage_Analysis

## Acknowledgments

We thank the Schmidt Ocean Institute, the captain, crew and pilots of and R/V Falkor (ROV *Ropos*) cruise FK160407 and R/V Thompson (ROV *Jason II)* cruise TN235, as well as the Kingdom of Tonga for allowing us to sample in their waters. We also thank P. Girguis for providing the 2009 samples, J. Becker for computational assistance, and C. Breusing for generating the metagenomically assembled genomes used in this study.

## Authors contributions

M.A.H., R.A.B., K.A. designed the study. M.A.H. performed the bioinformatic analyses. M.A.H. drafted the manuscript. M.A.H., R.A.B., K.A, K.M.K., and M.V.L. edited, reviewed, and approved the text and provided intellectual contributions.

