## Supplementary Figure 1a for "Phage interactions may contribute to the population structure and dynamics of hydrothermal vent microbial symbionts"

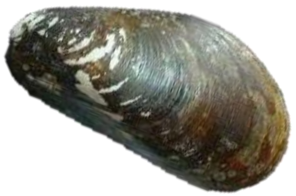

*Bathymodiolus Septemdierum*
98 metagenomes

0.005 ST phage/Mbp in metagenome

92 *Ca*. T endoseptemdiera symbiont MAGs

65 unique ST phage species clusters

7.67 CRISPR spacers/Mbp

0 prophage

59 ST singletons

6 ST groups

$\phi$ 25 → E
$\phi$ 28 → NE
$\phi$ 30 → E
$\phi$ 33 → E
$\phi$ 37 → E
$\phi$ 45 → E
$\phi$ 76 → E
