## Supplementary figures and images for "Phage interactions may contribute to the population structure and dynamics of hydrothermal vent microbial symbionts"

### Supplementary Figure 1b

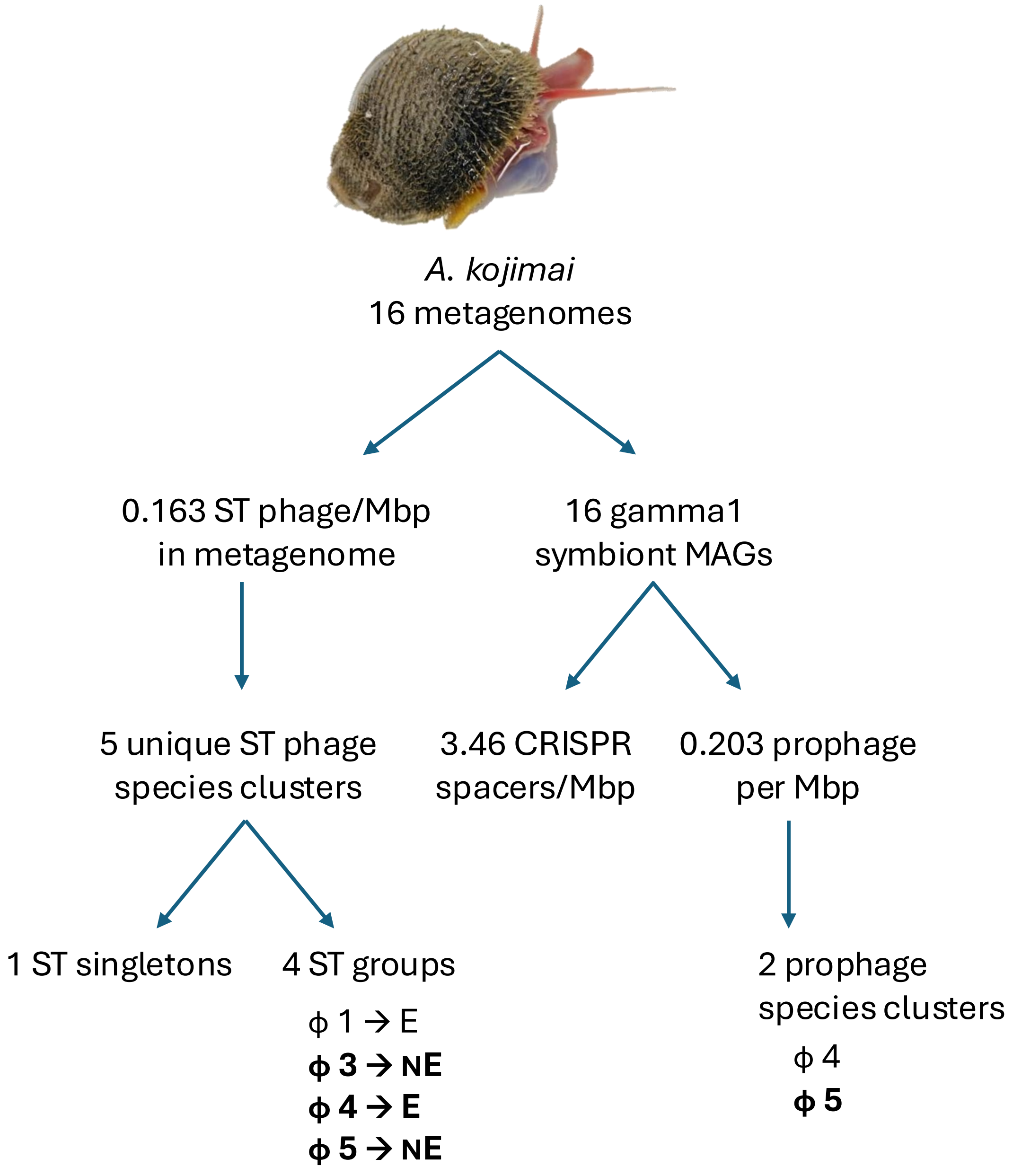

### Supplementary Figure 1c

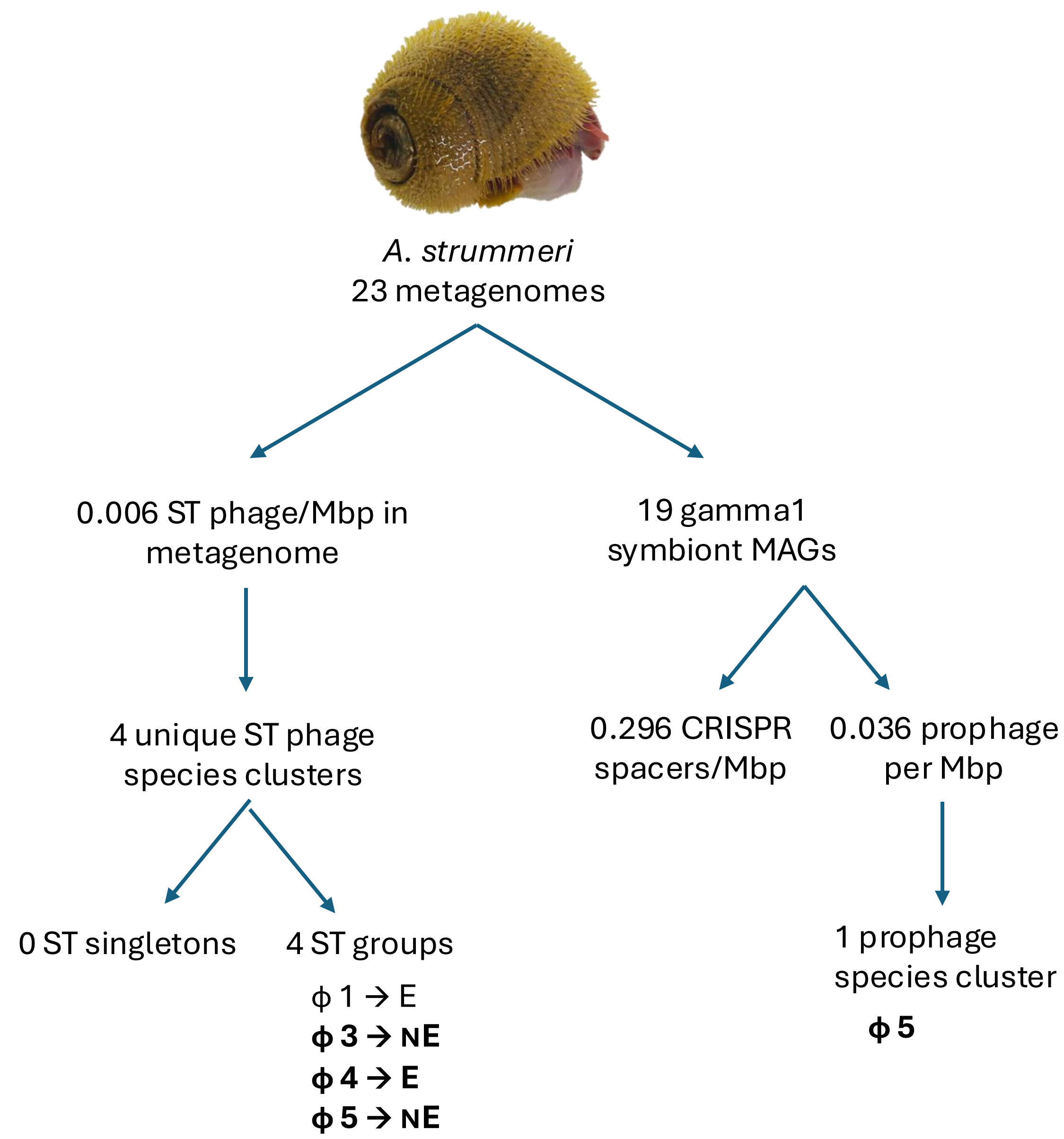

### Supplementary Figure 1e

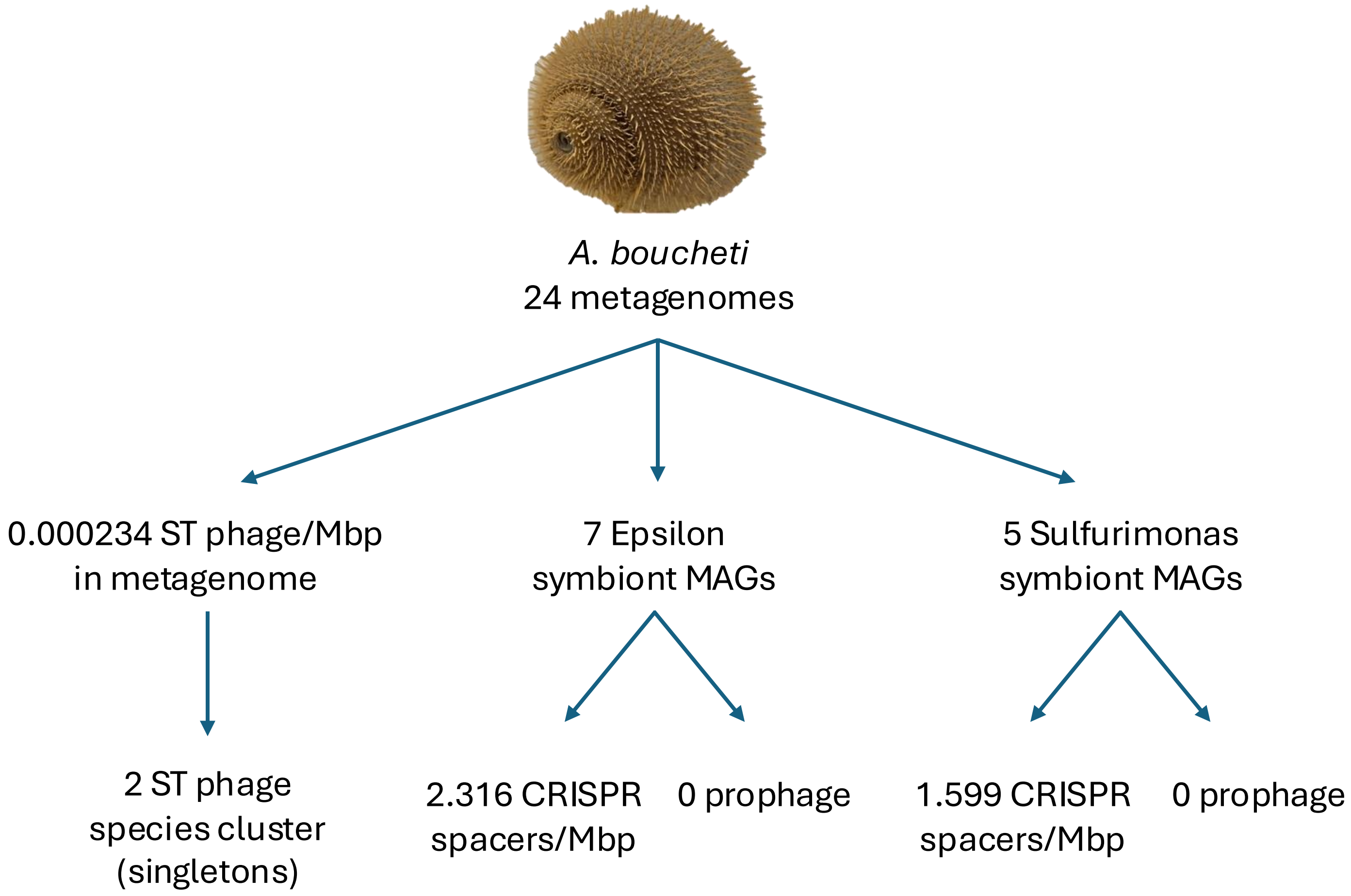

### Supplementary Figure 2

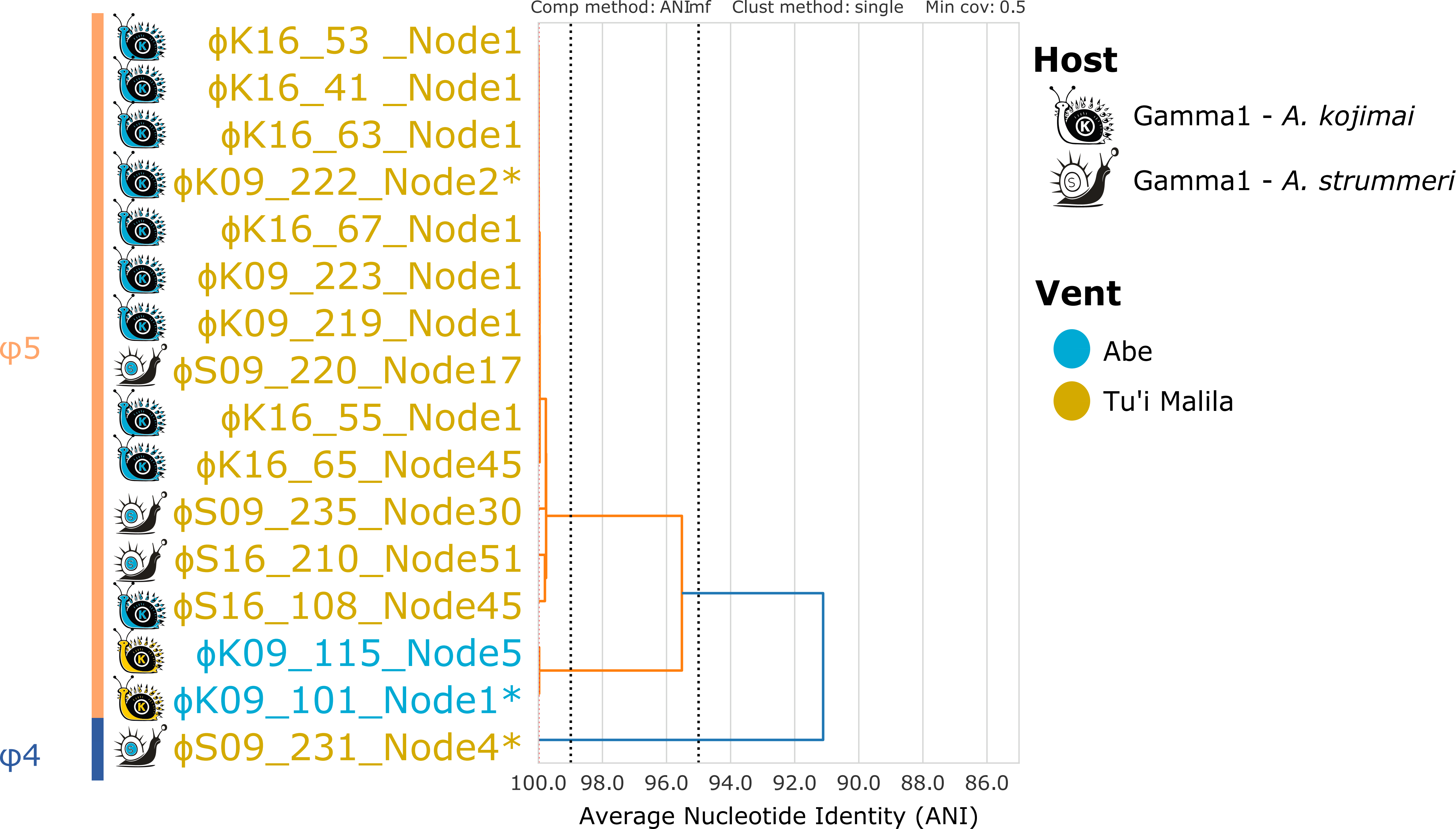
