## Supplementary Figure 1d for "Phage interactions may contribute to the population structure and dynamics of hydrothermal vent microbial symbionts"

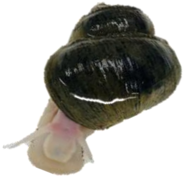

*I. nautilei*
58 metagenomes

0.003 ST phage/Mbp in metagenome

15 unique ST phage species clusters

9 ST singletons

6 ST groups

- ϕ 7 → NE
- ϕ 8 → NE
- ϕ 15 → E
- ϕ 16 → E
- ϕ 17 → NE
- ϕ 21 → NE
- ϕ 22 (MOX) → E

3 MOX symbiont MAGS

0 CRISPR spacers/Mbp

0 prophage

58 SOX symbiont MAGS

4.57 CRISPR spacers/Mbp

0 prophage
